# Deep ultraviolet light based wide-field fluorescence microscope for brain imaging

**DOI:** 10.1101/2020.10.27.342345

**Authors:** Deepa Kasaragod, Meina Zhu, Haruhi Terai, Koichi Kawakami, Hidenori Aizawa

## Abstract

Three-dimensional (3D) cellular scale imaging techniques that improve our understanding of the brain architecture is crucial for structural and functional integration and analysis of normal and pathological conditions in the brain. We have developed a wide-field fluorescent microscope using deep ultra violet (DUV) light emitting diode as the illumination source. This microscope employs oblique illumination of deep UV light and the optical sectioning is obtained on the tissue surface over a few micron thickness; largely attributed to the large absorption and hence low tissue penetration of the DUV light. Fluorescence emissions in the visible range that are spectrally separates allows for multiple channel of fluorophore detection using single or a combination of dyes. The fluorescence signal is captured using water immersion objectives and a color camera. Arduino Mega 2560 controlled 3-axis motorised microscope stage is developed for obtaining the wide-field imaging. To enable 3D imaging, the microscope setup is integrated with vibrating microtome that can slice thin sections for serial block-face imaging. In this paper, we show the versatility of the DUV microscope as a microscopy tool for use in neuroscience labs for a range of 2D brain imaging applications. We also present its applicability for 3D imaging of rodent brain in combination with whole brain staining protocol.

## Introduction

Three-dimensional (3D) cellular scale imaging techniques that improve our understanding of the brain architecture is crucial for the structural and functional integration and analysis of the normal and the pathological conditions. Recently there has been an surge of interest in 3D imaging of the brain involving the fluorescence microscope as a bridge to link the microscopic and macroscopic information. For example, habenula is a small brain region that is of specific interest pertaining to its link in motivation and decision making, depression and addiction. In animal models of chronic stress and depression, whether the structural changes can be seen in relation to the functional modifications, is still very much an open question and thus the volumetric analysis with cellular resolution of the heterogeneous substructures of habenula is of interest^1,2^.

Whole brain imaging has been realized by combining the different approaches of whole brain staining and optical clearing that allows for imaging of thick block of tissue sample using the optical sectioning ability of confocal, multi-photon, and light-sheet microscopy. However, all these microscopy setup are complex and require a great deal of optical expertise and expense to set up and utilise for 3D imaging, which do not make them a convenient choice for most of the laboratories to own.

Recently, a new method using deep ultraviolet light known as microscope using ultraviolet surface excitation (MUSE) was developed especially for imaging pathological samples in 2D^3^. This microscope makes use of two aspects of DUV light: 1. Large Stokes shift in the visible spectrum is seen with most of the conventional fluorophores when excited with DUV light 2. Strong absorbance of the DUV light and hence low penetration into the tissues is attained compared to the other spectrum of light. Owing to the low penetration depth, the optical sectioning ability of the imaging set up with the DUV light provides reasonable contrast to be used for imaging thick block of samples using a simple epifluorescence setup with the illumination light in oblique incidence. Simple off the shelf components have been used to develop microscopy systems for imaging of pathological specimens using DUV light^4–6^. The optical sectioning ability of the DUV light could be further improved through the use of water immersion objective^7^. Guo *et al*., showed the application of this concept to prototype 3D imaging based on single field of view imaging over depth for the visualization of mouse cerebral cortex^8^.

In this paper, we develop the DUV fluorescence microscope for wide-field imaging specifically for 2D/3D imaging of the brain with cellular scale resolution. We show the versatility of DUV fluorescence microscope that requires only a single wavelength light source and simple optical imaging set-p for a range of applications in brain imaging including cytoarchitecture imaging with cell nuclear staining, Nissl body imaging, immuno-fluorescence imaging, transgenic and viral vector transduction based fluorescence imaging. The applicability of this microscope for wide-field and cellular resolution serial block-face imaging of whole block of brain samples (habenula) is shown with whole brain staining of cell nuclear staining with Hoechst 33258. It is expected that this simplified microscope would be established as a useful tool for 3D microscopy for micro-macro analysis in neuroscience applications.

## Materials and Methods

### DUV Light Penetration and Fluorophore Excitation

The depth of light penetration in the biological tissues in DUV spectrum is limited owing to the strong absorption of the light by the biomolecules. This light penetration could be further reduced by using oblique incident of light illumination as well as with the use of the water immersion or oil immersion objective that increases the light angle of refraction in the tissue^7^. A schematic illustration of the depth of penetration using different schemes of UV light incidence in comparison to other wavelengths in the spectra are shown in Figure 1 (a).

**Figure 1.**
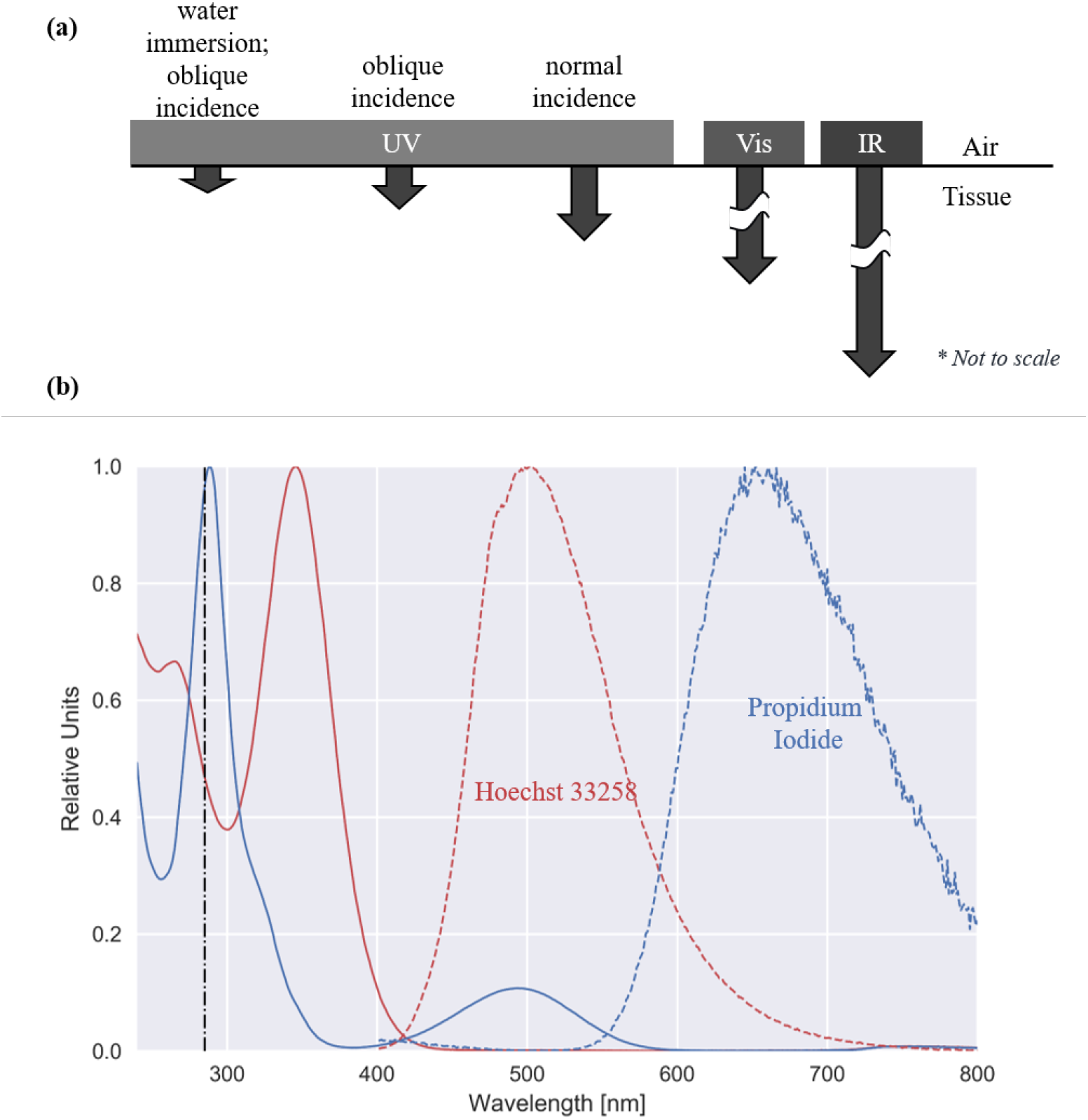
(a) Schematic showing the depth penetration of the DUV light in comparison to the wavelength in other spectrum as well as to oblique incidence illumination and water immersion based imaging configuration of DUV microscope. (b) The absorption spectra (solid lines) over the wavelength range of 240 to 800 nm and the emission spectra (dotted lines) obtained for excitation at 285 nm. The black dotted vertical line shows the excitation wavelength. The spectra were obtained using Varioskan LUX multimode microplate reader. The spectra are normalized with respect to peak of each curve.

DUV light is also well known for fluorescence of native biomolecules including NADH, tryptophan, collagen, etc. For example, enhanced green fluorescent proteins (eGFP) has a absorption peak at 280 nm owing to the presence of the aromatic amino acid residues in its chemical structure^9^. The native fluorescence of biomolecules as an improved contrast for brain imaging is seen in one of the channel of the multi-spectral image acquisition using color camera using the proposed microscope^10^.

A wide range of exogenous dyes that are conventionally used for brain imaging-for example: for cell nuclear imaging, neuronal Nissl body imaging, protein and antibody imaging are excitable at DUV wavelength in the range of 260-290 nm. With DUV excitation most of the conventional exogenous dyes show long Stokes shift and emission in the visible spectra. The emission spectrum for excitation at 285 nm and the absorption spectrum in 240-800 nm of some of the nuclear dyes (Hoechst 33258 and Propidium iodide) as measured using Varioskan LUX multimode microplate reader are shown in Figure 1 (b). The measurements were carried out using UV light transparent 96-well plate (Greiner UV-Star) with 1:50 dilution of the dye in water from Milli-Q water purification system (Millipore, Bedford, MA, USA).

### Wide-field Imaging Set-Up

The optical design of the imaging setup with the DUV light is essentially simple requiring use of no optical filters for emission or excitation. The optical design is similar to that of any conventional epifluorescence microscope but with oblique incident of light illumination onto the specimen. The DUV incident light excites the fluorophore in the visible spectrum and the images are acquired with single or multiple dyes that are spectrally separated using a color camera.

The current version of the DUV microscope uses DUV light emitting diode (LED) at wavelength 280 nm (Quark Technology, Okayama, Japan) with power incident on the sample as approximately 8.13 mW. The DUV microscope in the original built version used light source at 285 nm (Thorlabs, Japan) with incident power on the sample as approximately 1.25mW. The LED light is focused onto the surface of the sample in oblique angle of incidence. The images shown in the results sections are labelled DUV-280 and DUV-285 to highlight the light source used for imaging. The imaging was carried out using either the 10X and 20X water immersion objectives (Olympus UMPLFLN 10XW, Olympus UMPLFLN 20XW) with a working distance of 3.5 mm at 30 frames per second with a 12.3 MP color camera (Point Grey; GS3-U3-123S6C-C; 4,096 × 3,000 pixels) (Figure 2(a)). The field of view with the 10X objective was approximately 1.3 mm x 1.0 mm. The optical sectioning is achieved using DUV based fluorescence excitation limited to the tissue surface. The imaging setup is less complex compared to conventional fluorescent microscope setup because no additional blocking filters are required as the objective is opaque to DUV. The optical resolution of the imaging setup is obtained using USAF-1951 resolution target (2” x 2” Negative, 1951 USAF Hi-Resolution Target, Edmund Optics, Japan). Figure 2(b-c) shows resolving power up to Group 9 Element 3 (0.78 *μ*m) with the DUV microscope with 280 nm wavelength used as illumination light and 10X water immersion objective used for imaging.

**Figure 2.**
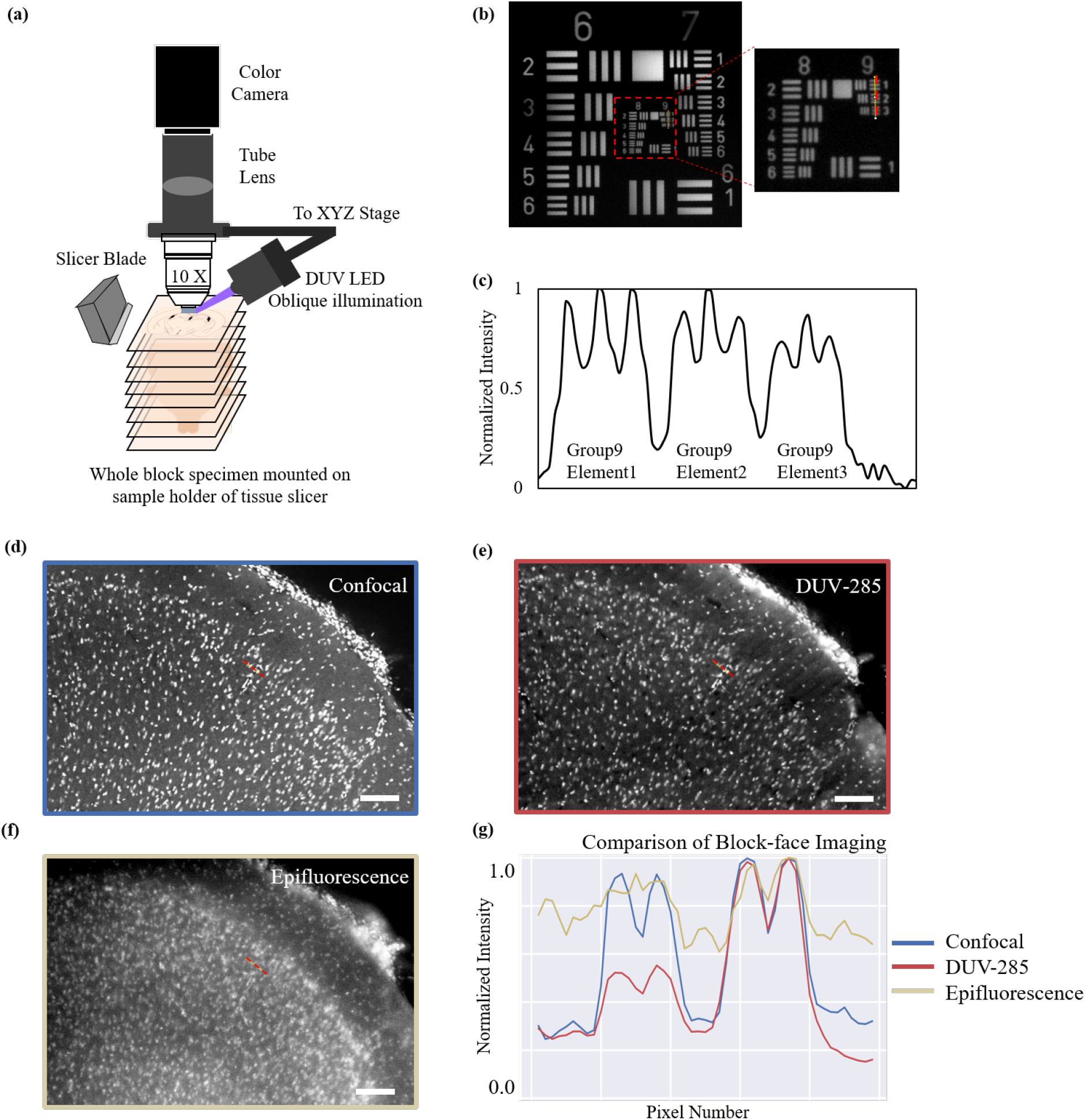
(a) Schematic of the 3D DUV imaging setup integrated with the tissue slicer for serial wide-field block-face imaging. (b) Image of a USAF-1951 resolution target showing resolving power up to Group 9 Element 3 (0.78 *μ*m) (c) The normalized intensity plot for dotted red lines over Group 9 of USAF-1951 resolution target. Block-face imaging of a thick block of mouse brain sample whole brained stained using Hoechst 33258 over exposed region on anterior cortical region using (d) Confocal microscope (Olympus FLUOVIEW FV1200MPE) (e) DUV microscope with illumination wavelength of 285 nm - blue channel of the color image (f) Epifluorescence microscope using visible light (405 nm) for excitation - blue channel of the color image (g) Intensity plot over the dotted red lines across each of the three images obtained from different microscopes show that the DUV performs better than conventional epifluorescence microscope for block-face imaging over thick tissue sample with slightly lower contrast obtained than that seen with the confocal microscopy system. Scale bar = 100 *μ*m.

**Figure 3.**
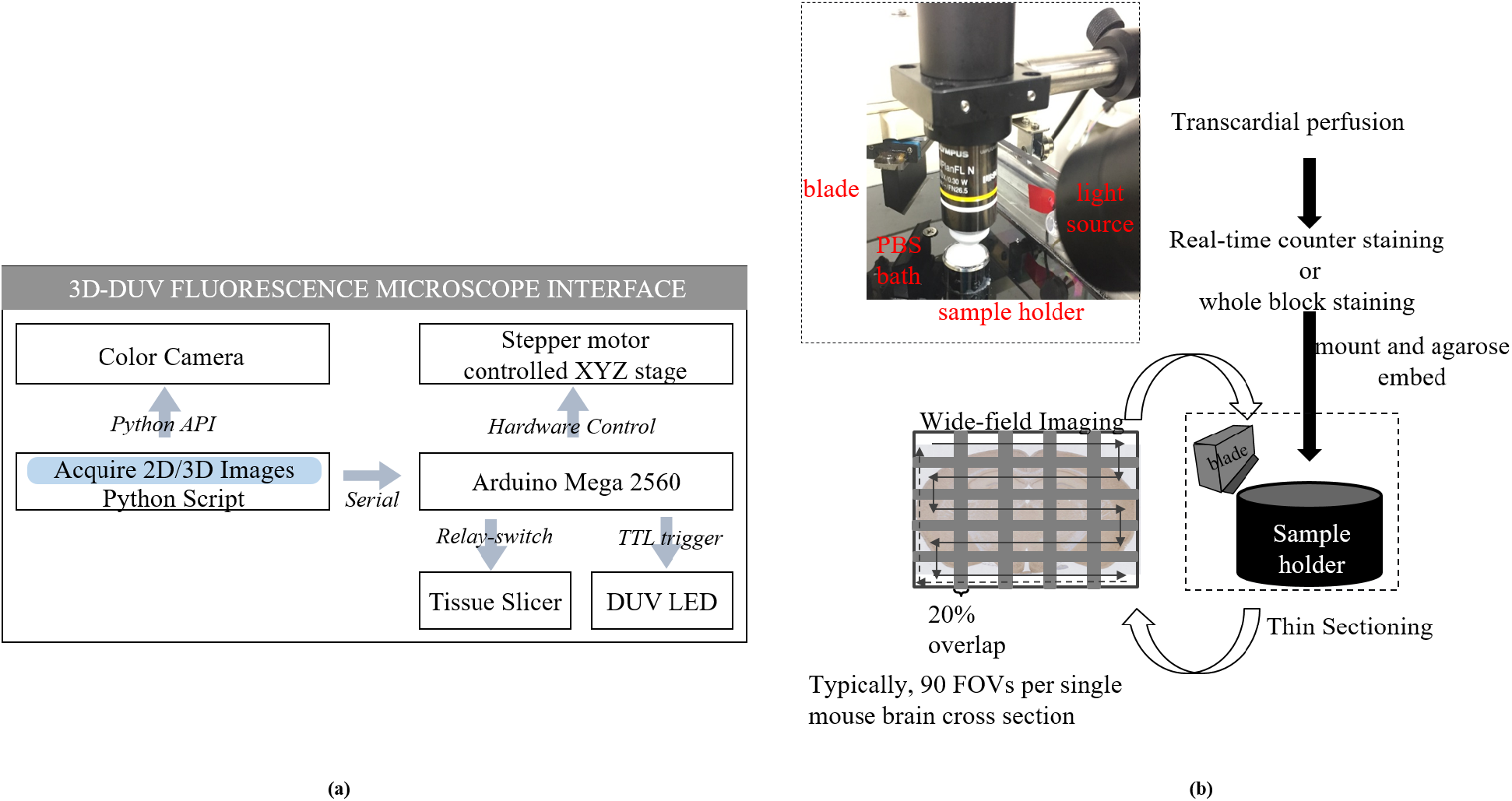
(a) A schematic of the hardware and software interface developed for 3D imaging using the DUV microscope. (b) Inset photograph of the imaging head showing the water immersion objective and the illumination source with the sample holder of the tissue slicer. Schematic showing the processes involved in the tissue preparation and image acquisition for serial wide-field block-face imaging.

The optical sectioning ability of the DUV microscope for imaging a thick block of tissue specimen is demonstrated using a whole brain Hoechst 33258-stained mouse brain sample. The whole brain staining protocol used is described in Section:Tissue Preparation and Staining Protocol later in the manuscript. The whole brain sample is sectioned coronally to expose an anterior cortical region for imaging. To compare the block-face imaging ability of DUV microscope, the same sample is imaged using three different types of fluorescent microscopes - 1. DUV microscope that excites at 285 nm; 2. Visible light Epi-fluorescent microscope at 405 nm with oblique incident similar to the DUV setup but with the relevant optical filters (Emission filter - FF01-446/523/600/677-25, Semrock; Excitation filter - FF01-390/482/563/640-25, Semrock) used; 3. Confocal microscope - Olympus FLUOVIEW FV1200MPE that used excitation light at 405 nm for Hoechst 33258. A quartz coverslip (Labo-USQ, 20 mm × 20 mm × 0.5 mm thickness, Daiko Seisakusho, Kyoto, Japan) is used to cover the tissue surface of the tissue block for imaging. Figure 2(d-f) shows the block-face images obtained using confocal, DUV and epifluorescence microscopy setup, respectively. Figure 2 (g) shows the normalized intensity profile over the dotted lines in Figure 2 (d-f). The dynamic range of the images were adjusted to obtain the similar noise floor outside the tissue region for all the three images and then normalization was carried out with respect to maximum grayscale values seen in each of the images. Blue channel of Hoechst 33258 is used for this analysis for DUV-285 and epifluorescence setup. This plot highlights the better performance of DUV microscope for block-face imaging in comparison to the epifluorescence microscope. DUV performance is on par with the contrast obtained with confocal microscopy setup for block-face imaging.

#### Motorised Microscope Stage

A customised motorised stage is designed for wide-field imaging which is a modified version based on the open-source based stage controller developed previously^11^. A flexible shaft is connected to 500 *μ*m/revolution micrometer actuated linear translator. Custom fabricated couplers are used to connect the shaft to the stepper motors (PK243M-01B; Oriental Motors, Japan) and the micrometers of the translation stage. PlayStation3 controller is used for manual control. Automated control of the stage is done using serial communication with Arduino Mega 2560. The master control is the custom written software using Python3 which communicates with the Arduino using the serial communication. For the stepper motor step size of 0.9*°*, a minimum resolution of 1.25 *μ*m is obtained with the translators.

#### 2D/3D Data Acquisition and Image Processing Pipeline

For 3D imaging, the tissue sample is embedded in the sample holder of the vibrating microtome (Compresstome VF-700-OZ-120, Precisionary Instruments). 2% agarose is used for the embedding the sample onto the holder. To allow fast cooling of the agarose, the bath surrounding the sample holder is filled with chilled PBS solution kept at 4 *°*C. The agarose is allowed to set for around 5 minutes before starting the serial slicing to remove the extra agarose layers on top of the sample. The desired thickness for serial sectioning is gradually arrived; starting with larger values of tissue thickness sectioning the top layers of agarose and gradually reducing the thickness with a step size of 5 *μ*m. Upon exposure of the tissue block-face, images per FOV are acquired as per the averaging and Z-stack values input for all the field of views (FOVs) across the entire wide-field imaging. Usually, four images are acquired per FOV for averaging over one Z-position in focus. The number of X and Y translations required to cover the entire tissue section or tissue section of interest is set based on the extent of the tissue. The image acquisition and stage translation are timed asynchronously and empirically. The XYZ stage that holds the imaging unit is brought back to the original location after completion of an entire wide-field image acquisition of the exposed section. Microtome slicer is controlled using a relay switch through Arduino Mega 2560 to automatically start the sectioning of the tissue after completion of the wide-field data of the previous tissue section. To allow the slicer blade to slice off the next section, the imaging head is translated 5mm away from the original position. The slicer then sections the exposed block face at the desired thickness followed by returning of the blade back to the original position. The imaging head is then moved back to the original position and the the whole process is repeated till the imaging of the entire block of tissue sample is carried out over serial sections of desired thickness.

For 2D imaging, quartz coverslip is used on top of the sample and wide-field mosaics are obtained without the use of tissue slicer in the same imaging setup.

The parameters corresponding to each of the imaging session is described using YAML files, which is a common plain text human readable serialization language^12,13^. This YAML file is utilized in the image processing pipeline for automated processing of each brain sections. The YAML files can also be opened and edited using any standard text editor. An example of the YAML file used in imaging session of DUV microscope is shown in Figure 4.

**Figure 4.**
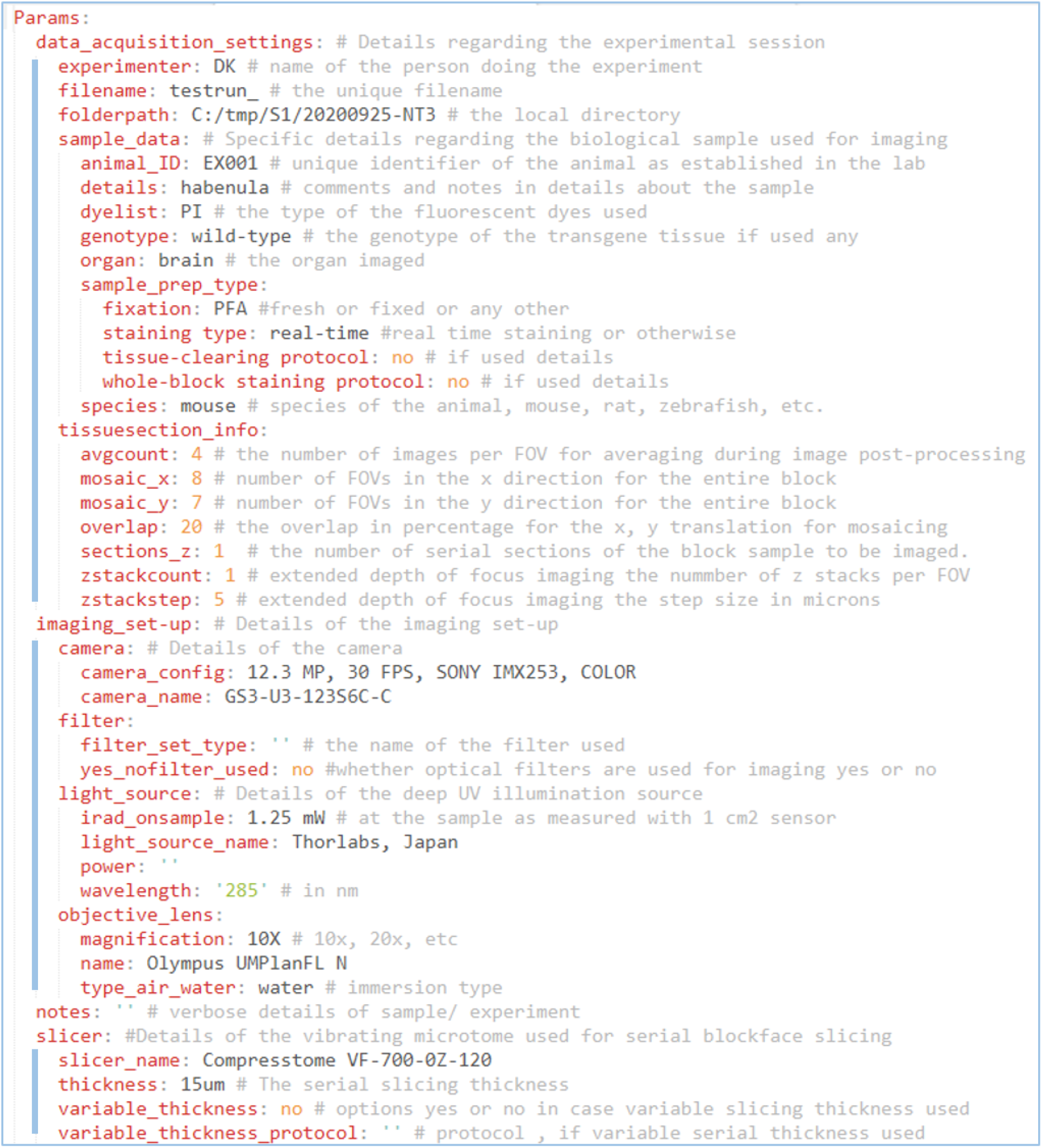
An example configuration file in YAML. The metadata pertaining to parameters corresponding to each of the experimental session is listed and described using human-readable plain text YAML file. This YAML file is utilized in the image processing pipeline for post-processing of wide-field images.

The camera is controlled by the FlyCapture2 API. The image acquisition and image analysis is carried out using custom written software using Python3. The image processing steps include color correction, flat-field correction and average over four images per FOV. Extended depth of focus imaging is optionally carried out if multiple z-stack images are acquired for each FOV. Wide-field mosaicing is obtained using Microsoft Image Composite Editor for each of the brain sections. 3D reconstruction is carried out using ImageJ plugin.

### Tissue Preparation and Staining Protocols

For 2D imaging, the coronal brain sections of 50 *μ*m thickness or thick tissue blocks without serial sectioning were used with quartz coverslip. The animals used in this study for brain imaging are wild-type C57BL/6J strain mouse, unless specified otherwise. Two different lines of zebrafish were used : SAGFF31B transgenic line14 and wild-type strain. All the animal experiments, care and handling procedures were performed in accordance with the guidelines provided by the Animal Care and Use Committee of Hiroshima University.

The tissue preparation and staining protocols of the different fluorescent imaging applications shown in this study using DUV microscope was carried out as following:

1. Cell type specific imaging using fluorescent markers: dopaminergic neurons were specifically labelled using the anti-Tyrosine Hydroxylase (TH) antibody (1:250, rabbit polyclonal, AB152, Sigma Aldrich). This was done on 50 *μ*m thick coronal sections in the midbrain area specifically for the neurons in the ventral tegmental area (VTA) and the substantia nigra. The antibody staining protocol is as follows: The slices are washed in 1X PBS thrice for 10 min each at room temperature (RT) on a shaker. The slices are then incubated in 0.5% Triton X-100 in 1X PBS for 2 hours at RT on a shaker. Primary antibodies are added to the existing solution and is incubated overnight at 4*°*C. The following day the slices are washed thrice for 10 min each in 1X PBS at RT on a shaker. The slices are incubated with secondary antibody (goat anti-rabbit IgG H&L ; Alexa Fluor^®^ 594; ab150080, abcam) for 2 hours and then washed thrice in PBS and mounted on glass slide with quartz coverslip.
2. Recombinant adeno-associated virus (AAV) vector expression imaging of enhanced green fluorescent proteins (eGFP): Stereotaxic injection of AAV8-CAG-GFP (Addgene ID: 37825-AAV8 https://www.addgene.org/37825/) on 11 weeks old male mouse. Injection coordinates were −1.75 mm posterior from Bregma, 0.5 mm bilateral to the midline and 2.8 mm in depth from the surface. Imaging was done 9-month after the viral vector injection.
3. Nissl body imaging using fluorescent dyes: For Nissl body staining NeuroTrace 530/615 Red Fluorescent Nissl Stain (N21482, Invitrogen, USA) as well as Propidium iodide (P1304MP, Invitrogen, USA) were used with the coronal sections for staining with 1: 250 and 1:500 dilution, respectively in PBS.
4. Transgenic expression of the fluorescent biomarkers: The imaging of the transgenic fluorescent reporter was shown on the zebrafish using SAGFF31B^14^. 50 *μ*m coronal sections were imaged using laser scanning confocal microscope (Olympus BX51WI with Yokogawa CSU-X spinning disk confocal scanner unit and Andor iXon Ultra EMCCD) as well as DUV microscope for GFP fluorescent expression.
5. Neuronal pathway imaging: The habenulo-interpeduncular projection in zebrafish was visualized using lipophilic tracer DiI (D-3911, Invitrogen, USA). DiI granule was applied to the right and left habenula by placing on the tip of a sharp glass needle followed by incubation at 37*°*C for 48 hours to allow the dye diffusion to occur^15,16^.
6. Whole Brain Staining: For whole brain staining using the nuclear staining dyes, the osmotic shock protocol based on varying the sucrose amount was used^17,18^. Mice were anesthetized with an intraperitoneal injection of ketamine and were transcardially perfused with PBS followed by 4% paraformaldehyde (PFA) dissolved in PBS. Excised brains were post-fixed in 4% PFA in PBS overnight. Next day the brains were washed in PBS twice for 2 hours. The mouse brains were stained with Hoechst 33258 dye (Dojindo Laboratories, Kumamoto, Japan) or Propidium iodide as follows: The brains were immersed in staining solution (PBS containing 10mg/ml Hoechst 33258 or Propidium iodide, 0.1% TritonX-100, and w/v sucrose: Day1 - 10%; Day2 - 20%; Day3 to Day7 - 30%; and Day8 - 0%) at 55*°*C with gentle shaking.
7. Real-time staining: For real-time staining, the rodent brains were extracted after transcardial perfusion and PFA fixed overnight. Next day, the sample is washed twice in PBS is stored in PBS at 4*°*C until use. For imaging, the sample is embedded in the microtome sample holder using agarose. The chill bath used for solidifying the agarose is replaced with the diluted nuclear dye like Propidium iodide, etc in PBS (Results not shown).

## Results

### DUV fluorescence microscopy for brain imaging

Along with the DUV microscope, 2D imaging is also performed using conventional epifluorescence microscope (Olympus MVX10) on mouse brain sections and using scanning laser confocal microscope (Olympus BX51WI with Yokogawa CSU-X spinning disk confocal scanner unit and Andor iXon Ultra EMCCD) for zebrafish samples.

Figure 5(a) shows the schematic of the mouse coronal section showing the bilateral injection site of the AAV. Figure 5(b) shows eGFP expression in lateral habenula (dotted region) after the injection of the AAV vector with the fluorescent protein (green color in DUV RGB image). The blue color in the image is from the background tissue autofluorescence which highlights the anatomical landmark especially with respect to the medial habenula. The imaging was done on a thick sample and so no epifluorescence image is shown alongside.

**Figure 5.**
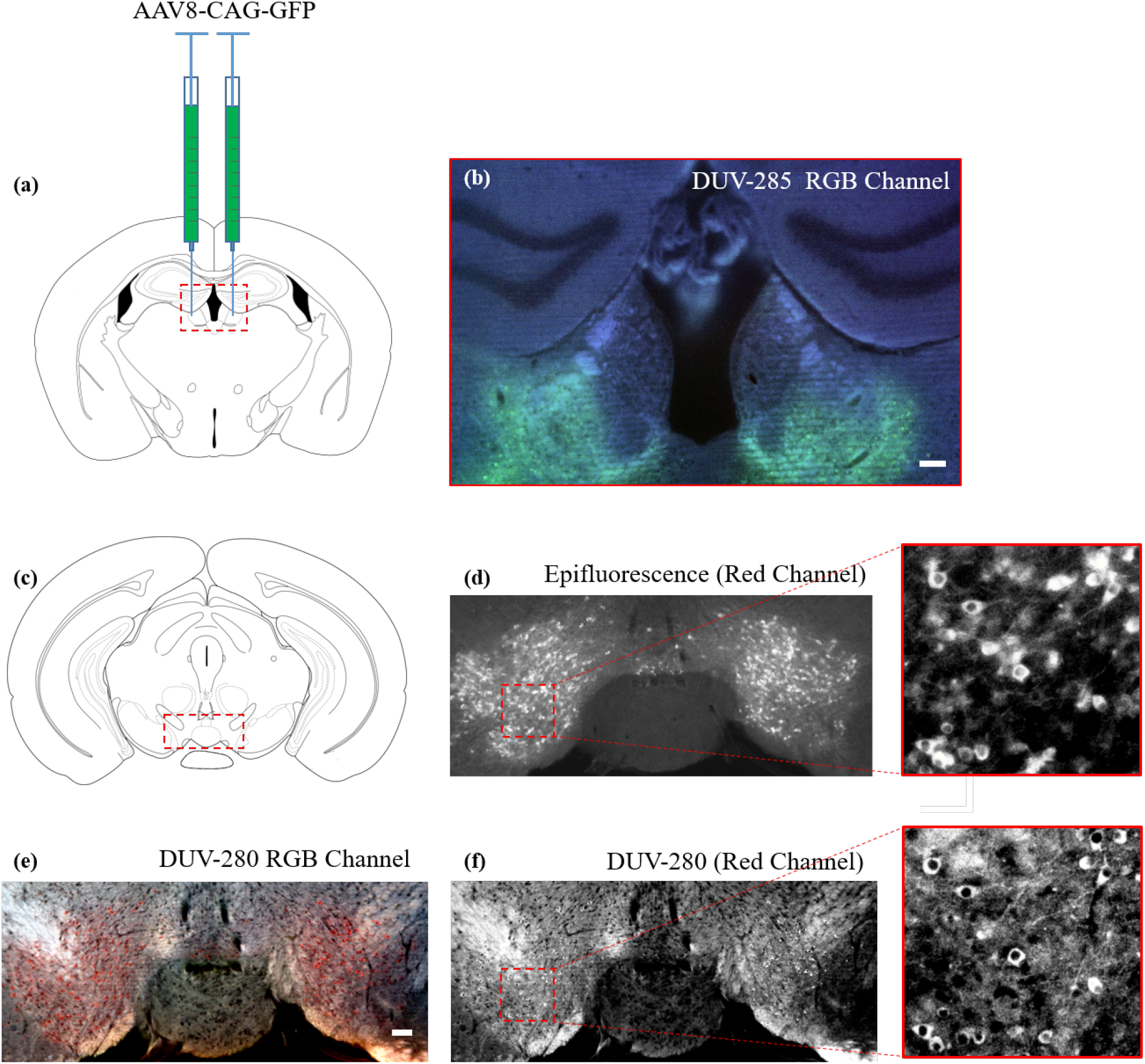
(a) Schematic of the mouse coronal section showing the AAV injection site with RGB image of the GFP expression in the AAV injected site obtained using DUV microscope for red dotted region shown in (b). (c) Schematic of the coronal section used for cell-specific imaging of dopaminergic neurons in the VTA of the midbrain labelled using anti-TH antibody with red fluorescent AlexaFluor 594. Red dotted region is shown (d) Red channel image obtained from epifluorescence microscope (e) Color image obtained from DUV microscope (f) Red channel extracted from the color image of DUV microscope. Scale bar = 100 *μ*m.

Figure 5(c) shows the schematic of the coronal section of the midbrain area used for anti-TH immunofluorescence imaging. Figure 5(d) shows the epifluorescence image in grayscale (e) shows the DUV RGB image and (f) shows the DUV red channel image in grayscale for the dotted red regions highlighted in Figure 5(c). Zoomed in ROI from epifluorescence and DUV are also shown side by side for a better comparison of the better contrast and sharp image quality obtained with DUV microscope albeit the presence of white matter autofluorescence.

Figure 6(a-e) shows the DUV microscope for imaging the fluorescent Nissl staining using NeuroTrace 530/615 and Propidium iodide on the coronal sections containing the striatum. Images are shown as DUV RB channel images as a composite of red (Nissl) and blue (white matter autofluorescence) by removing the irrelevant green channel. For a comparison between the DUV microscope and conventional epifluorescence microscope, Figure 6(b-c) shows the DUV red channel and epilfuorescene in grayscale, respectively. The number of cells seen in DUV are less than that in conventional epifluorescence image owing to the surface excitation feature of DUV due to the DUV light penetration based optical sectioning.

**Figure 6.**
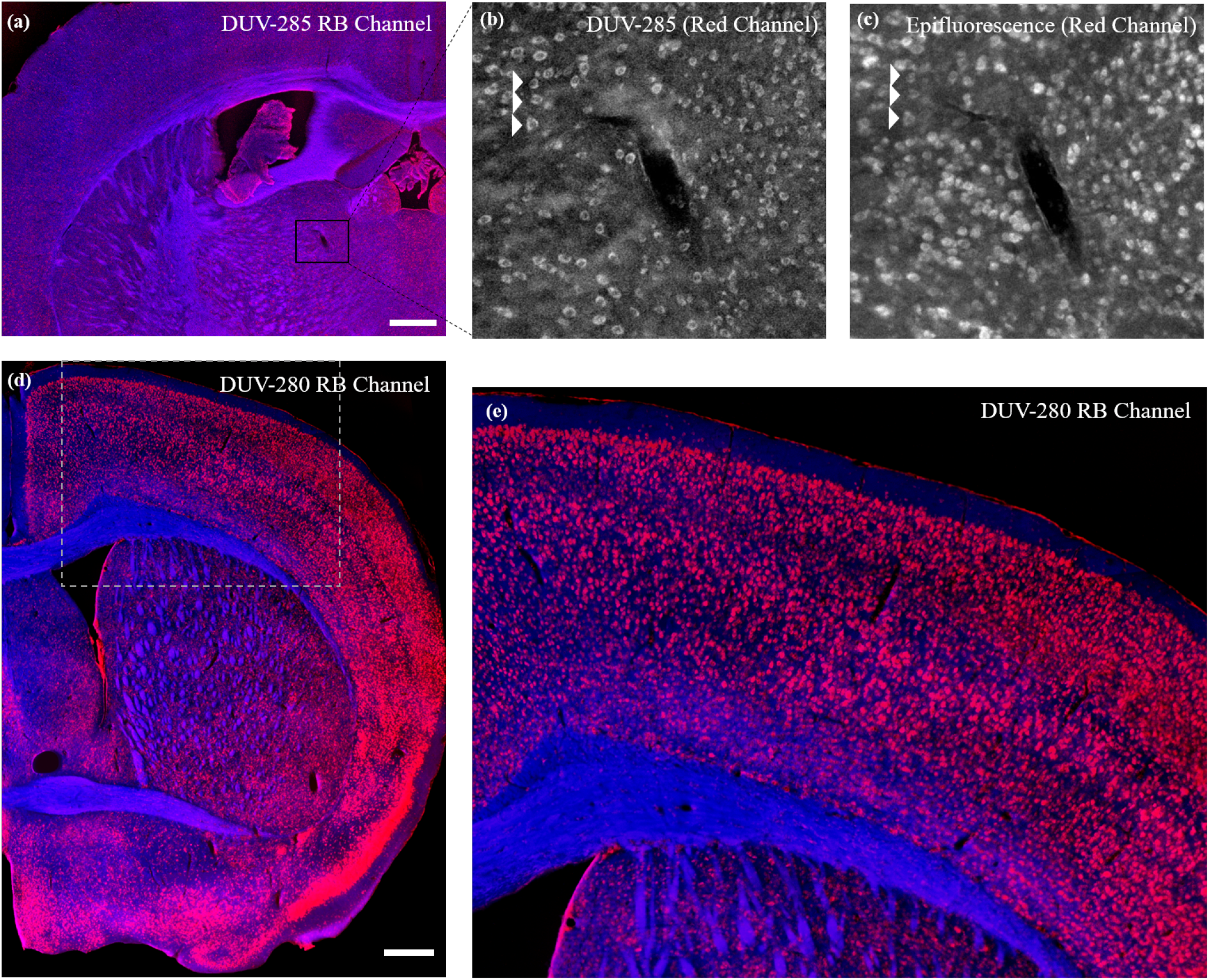
Red fluorescent Nissl body imaging using the DUV microscope with (a-c) NeuroTrace 530/615 (d-e) Propidium iodide staining on anterior coronal section of mouse brain comprising the striatal region. (a) Image obtained from DUV microscope showing only autofluorescence (blue) and Nissl (red) channels with zoomed region showing only the red channel in grayscale for (b) DUV microscope (c) epifluorescence microscope. Scale bar = 500 *μ*m.

DUV microscope is a not a contender to the light sheet microscope in terms of its advantages for zebrafish imaging. As DUV is a surface excitation based approach with limited depth penetration, the imaging applications of zebrafish would be limited to the ex-vivo imaging after the extraction of the brain samples. Nevertheless, the applicability of the DUV microscope for imaging the thin sections of zebrafish with transgene fluorescent expression and to study the neural pathways are shown in Figure 7. Figure 7 (a)-(c) are from the expression of the SAGFF31B transgene line, with Figure 7(a) showing the transverse section that was imaged. This image was taken with a scanning laser confocal microscope and was mosaiced over series of FOVs using ImageJ MosaicJ Plugin. Only the high saturated signals seen around the hypothalamus areas are visualized using DUV microscope (Figure 7(c)). One of the reasons why the green fluorescence expression other than the high intense signals around the hypothalamus is not observed with DUV microscope might be due to the low LED irradiance.

**Figure 7.**
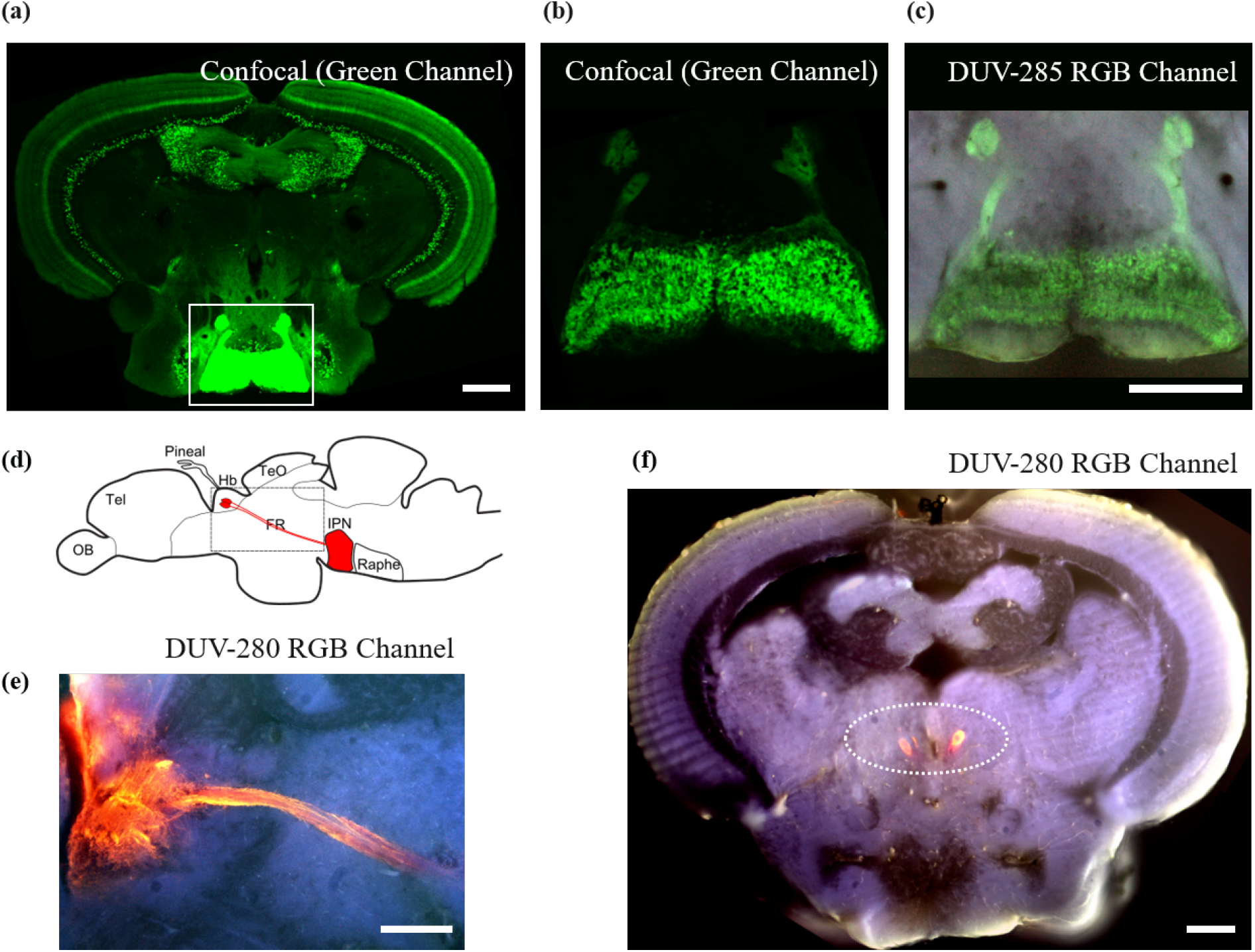
(a) Mosaic image as obtained from laser scanning confocal microscope showing GFP cells of SAGFF31B transgenic line zebrafish in one of the coronal cross section. Highlighted ROI shows the hypothalamic cells obtained with (b) Scanning confocal and (c) DUV microscope. (d) Schematic illustration of a lateral view of an adult zebrafish brain showing the habenular (Hb) projections to the ventral midbrain FR - fasciculus retroflexus; IPN - interpeduncular nucleus (e) Transverse sections of the bounded region shown in the schematic showing the habenular axons projecting towards ventral interpeduncular nucleus (f) Coronal section of another sample of zebrafish (wild-type) showing the fasciculus retroflexus stained red. Scale bar = 200 *μ*m.

Figure 7(d) shows the schematic to highlight the habenulo-interpeduncular projection seen in the zebrafish. Figure 7(e-f) shows the transverse and coronal sections obtained from two different zebrafish brain samples highlighting the habenulo-interpeduncular projection. The dotted circle in the coronal section in Figure 7(f) is drawn to highlight the fluorescence signal seen in fasciculus retroflexus cross-section. Also, notice the enhanced anatomical landmark contrast seen in the DUV color image from the background signal, which is often not seen in the conventional fluorescence microscopy of zebrafish.

### Serial wide-field block-face imaging of the rodent brain

As a prototypical approach, the 3D microscopy involving serial block-face imaging of whole brain nuclear stained brain sample is shown. This is illustrated with 3D reconstruction of the unilateral habenula structure over 10 serial sections of 15 *μ*m thickness each. A representative wide-field coronal section of the block-face imaged sample as DUV color image is shown in Figure 8(a). A close-up of the bilateral habenula is shown in Figure 8(b) along with the virtual histology like image obtained from the DUV color image. ImageJ VolumeViewer was used to create the 3D rendering of the unilateral habenula using the blue channel signal of the DUV image as seen in grayscale in Figure 8(c). Note that the blue channel also shows the autofluorescence signals more prominently seen in the medial habenula.

**Figure 8.**
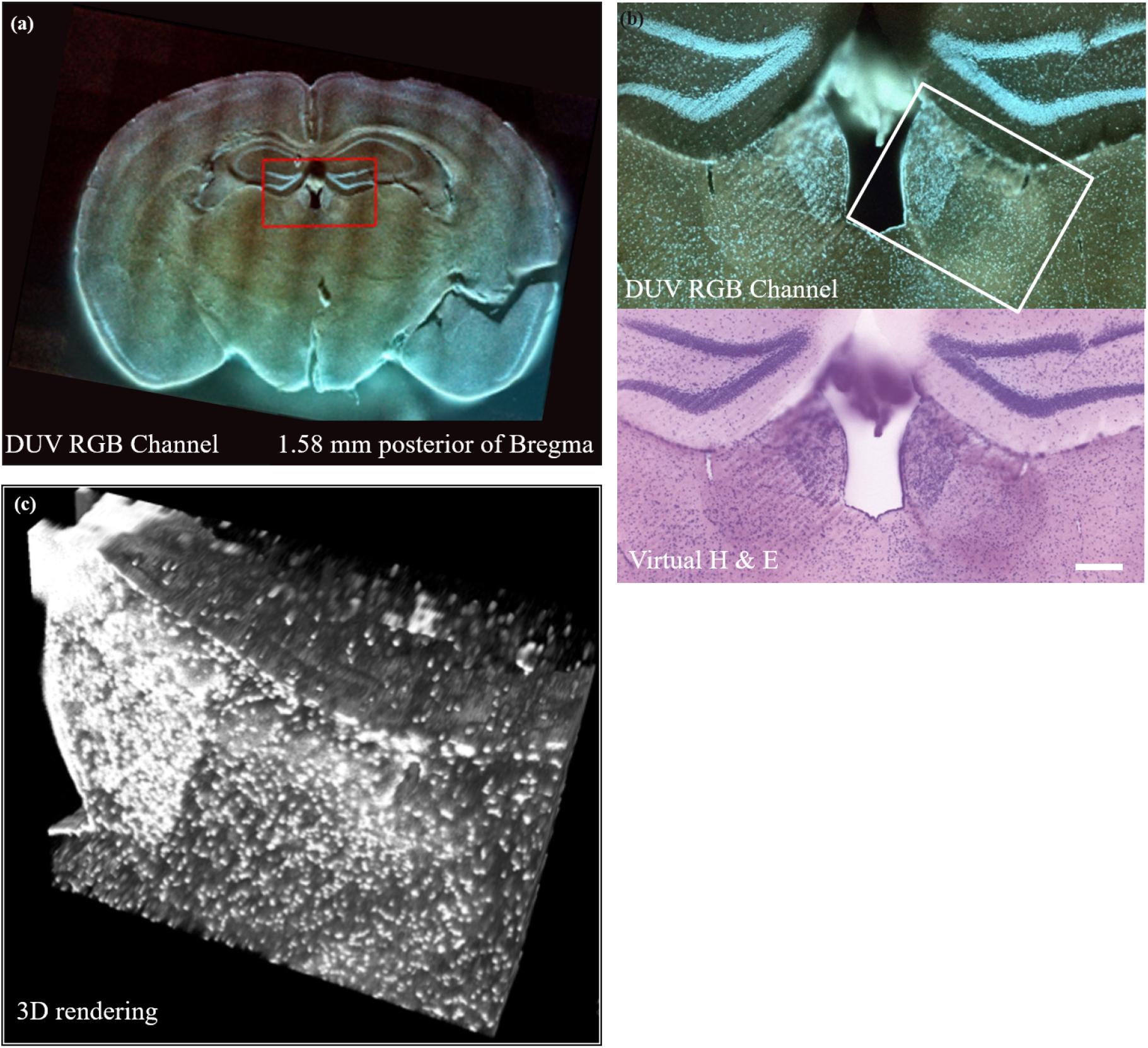
(a) Wide-field DUV RGB image of the coronal section of habenula 1.58 mm posterior of Bregma imaged using DUV microscope (b) Zoom in of red bounded box region showing bilateral habenula -DUV RGB image and virtual H & E stained image obtained from DUV image (c) Habenula 3D rendering of 10 stacks of 15 *μ*m serial sections. 3D rendering is carried out using only the blue channel of Hoechst 33258 signal. Scale bar = 200 *μ*m.

## Discussions

The underlying principles of DUV microscope are same as the recently developed microscopy with ultraviolet surface excitation (MUSE)^3, 7^. MUSE has been proposed as a rapid tool for obtaining histology like images for pathology labs. In this study, we are showing the possibility of extension of the utility of MUSE/DUV microscope as a alternate for conventional epifluorescence microscope for any neurobiology/neuroscience research labs for thin or thick section imaging. DUV microscope is presented here as a simple optical design that can be used as a replacement of the conventional microscope in research labs for both 2D and 3D microscopy.

DUV light is excitable for a range of fluorophores including conventional nuclear dyes like Hoechst 33258, Propidium iodide, Nissl fluorescent stains like Neurotrace 530/615, Propidium iodide, transgene expresssion of fluorescent proteins like GFP, eGFP, fluorescent secondary antibodies, etc. The feasibility of protein specific imaging using DUV for exciting Quantom dots have been recently shown^19^. The list of fluorophores, both endogeneous and exogeneous, shown here is not exhaustive and future studies are required to create the list of the dyes that are and can be engineered for excitation using DUV spectrum.

The serial block-face imaging ability of DUV microscope integrated with a tissue slicer was shown here using habenula 3d mapping Figure (8 (c)). The z-sectioning thickness is limited by the uniform cutting ability of the tissue slicer and the type of the tissue preparation involved. In this study, for whole brain stained sample, we were able to uniformly serially section to a thickness of 15 *μ*m. The serial sectioning thickness can be improved further by combining with microtome that can slice paraffin wax embedded tissue sections^8^. Our focus with this study was to develop the DUV microscope specifically for PFA fixed samples, without requiring extensive tissue embedding or preparation other than the staining protocols.

Optical microscopy conforms to the laws of diffraction and thus moving to the shorter wavelength of the DUV spectrum has the added advantage of the better spatial resolution under diffraction limited regime. Moving to the DUV spectrum also involves the significant presence of ’autofluorescence’ coming from the endogeneous UV excitable biomolecules attributed to excitation of the aromatic amino acid residues like tryptophan, tyrosine and phenylalanine^20^. Although these molecules are excitable in the range of 220-295 nm, their emissions are limited from 280 to 350 nm and outside of the visible spectrum of our interest. In the brain samples, white matter autofluorescence with DUV excitation is significant. Although in this paper we have not spectrally separated the white matter autofluorescence signal from the fluorophore emission of interest, custom trained algorithms for DUV images can be designed using supervised learning based algorithm as a future study to transform DUV color images into spectrally separated signals along with the white matter autofluorescence. The white matter autofluorescence also provides the anatomical landmark in the interpretation of the fluorescent images of interest.

## Conclusions and Future Perspective

We have developed a low-cost single wavelength based simplified optical design for wide-field serial block-face microscopy for 2D/3D brain imaging. We have shown its extensive utility for the conventional applications of 2D brain imaging. We have also shown how this microscope could be utilized for the 3D imaging of the brain in combination with a tissue slicer. As a future application for this microscope, multiplex protein imaging in combination with the whole brain optical clearing and staining technique shall be tried out.

## Acknowledgements

The authors acknowledge the discussions with Dr. Richard Levenson, Dr. Farzad Fereidouni and Mr. Austin Todd, University of California, Davis Medical Centre, Sacramento, CA, USA. We are grateful to the National BioResource Project (NBRP) of Zebrafish, Core Institution, for providing the SAGFF31B zebrafish transgenic lines. This study was supported by HIRAKU Consortium Start-up Grant 1818ZE7279, Female Researcher International Joint Research Grant 1811730, JSPS Kakenhi 19K206740A (to DK); JSPS Kakenhi 20HP8021, NBRP/Fundamental Technologies Upgrading Program from the Japan Agency for Medical Research and Development (AMED) (to KK) and 15K217150B International Collaboration Activity Support Grant, JSPS Kakenhi JP26112010, JP19H05723, JP17K19459 and a Grant-in-Aid for Integrated Research on Depression, Dementia and Development Disorders (18dm0107093h0003) (to HA).

## Author Contributions

DK and HA designed the instrumentation and the experiment(s); DK conducted all the imaging experiments and analysed all the results. ZM contributed to the whole brain staining experiment of mouse and performed the recombinant viral vector based experiments; KK generated SAGFF31B transgenic zebrafish; HT and HA performed the zebrafish breeding, maintenance and brain preparation; DK wrote the initial draft of the manuscript and all authors contributed to the modifications and revisions to the final draft. All authors reviewed the manuscript.

